# Utilizing full-length 16S rRNA sequencing to assess the impact of diet formulation and age on targeted gut microbiome colonization in laboratory and mass-reared Mediterranean fruit flies

**DOI:** 10.1101/2024.12.27.630527

**Authors:** Charles J. Mason, Rosalie C. Nelson, Mikinley Weaver, Tyler Simmonds, Scott Geib, Ikkei Shikano

## Abstract

Insect gut microbiomes have important roles in overall host health and how hosts function in the environment. In laboratory and mass-reared insects, gut microbiomes can differ in composition and function compared to wild conspecifics. For fruit flies, like the Mediterranean fruit fly (medfly; *Ceratitis capitata*), these changes can influence male performance and behavior.

Overall, understanding factors that influence the ability of bacteria to colonize hosts is an important for the establishment of lost or novel microbiota into mass-reared insects. The goal of this study was to evaluate how host age and diet inoculation method influenced bacterial establishment in laboratory and mass-reared medfly. We used an *Enterobacter* strain with antibiotic resistance and coupled it with full-length PacBio 16S rRNA sequencing to track the establishment of a specific isolates under different adult dietary conditions. We also used two longstanding reared lines of medfly in our study. Our results identified that diet had a strong interaction with age. Host medfly fed a liquid diet with the target bacteria were able to be colonized regardless of age, but those fed a slurry-based diet and separate water source were more resilient. This was consistent for both fly rearing lines used in the study. 16S rRNA sequencing corroborated the establishment of the specific strain, but also revealed some species/strain-level variation of *Enterobacter* sequences associated with the flies. Additionally, our study illustrates that long-read 16S rRNA sequencing may afford improved characterization of species- and strain-level distribution of Enterobacteriaceae in insects.

**Importance:** Insects form intimate relationships with gut microorganisms that can help facilitate several important roles. The goals of our study were to evaluate factors that influence microbial establishment in lines of the Mediterranean fruit fly (medfly), an important pest species throughout the world. Mass-reared insects for sterile insect technique often possess gut microbiomes that substantially differ from wild flies, which can impact their performance in pest control contexts. Here, we show that liquid-based formulations can be utilized to manipulate the gut microbiota of mass-reared medfly. Furthermore, using near full-length 16S rRNA metabarcoding sequencing, we uncovered strain-level diversity of that was not immediately obvious using other approaches. This is a notable finding, as it suggests that full-length 16S rRNA approaches can have marked improvements for some taxa compared to fewer hypervariable regions at approximately the same cost. Our results provide new avenues for exploring and interrogating medfly-microbiome interactions.

## Introduction

The insect gut microbiome is an important component of host health and environmental interactions (1–3). For many insects, the gut can harbor high microbial density (4), and beneficial partnerships may assist in aspects of digestion and nutrient assimilation (5, 6), detoxification (7, 8), and interactions with antagonistic microorganisms (9, 10). Robustness and fidelity of gut symbioses can differ between different insect lineages (3), meaning that some species may exhibit strong, host-microbiota feedbacks of individual taxa while others may exhibit more variation in what establishes and occupies the digestive system. Determining factors that affect microbial turnover and plasticity in insect gut systems can provide insight into the strategies of host microbiomes to resist or succumb to invasion, and how these processes ultimately impact host functioning. From a practical standpoint, introducing depleted, lost, or different strains to insect gut microbiomes to improve insect performance may be exploited for improved agricultural services by some insect species (11–14). In particular, reintroduction of bacterial strains lost or depleted in mass-rearing conditions has been proposed to improve the performance of mass-reared fruit flies (Diptera: Tephritidae) produced for sterile insect technique (SIT) (11, 15, 16).

Mass production of insects often involves derivations from natural conditions for peak efficiency of production. Some of these include changes to diet composition, population densities, and steps to limit undesirable microbiota can collectively impact the microbiome associated with these insects. Mass production can have determinantal impacts on individual performance parameters compared to wild populations, which has been the case for mass-reared fruit flies, including the Mediterranean fruit fly (medfly; *Ceratitis capitata*). Fruit fly microbial communities shift rapidly through the domestication from the wild flies initiating the colony (17–19). Recapitulation of bacteria from wild sources has been pointed to as a potential avenue for improving the competitiveness of sterilized male fruit flies (20–23).

Medfly larvae and adults vary in their gut microbiome composition and diversity across space and time (24, 25). Prior studies have reported that fruit fly gut microbial populations and communities can vary between fruits and diets (26–28), rearing sources (29, 30), developmental stages (31–33), and as adults age (34). Adult medfly microbial titers shift over short durations following adult eclosion (34, 35); expanding from low numbers (< 10^3^ colony forming units (CFUs) per fly) in colony and mass reared flies only to recover within 72h (> 10^7^ CFUs per fly) (Supplemental Figure 1) (34). Despite variability between individuals, there are generally similar taxonomic groups that comprise these communities such as *Klebsiella, Enterobacter*, *Enterococcus,* and *Pseudomonas* (15, 16, 36). How much strain-level diversity is present in fruit fly systems is currently unclear, since recent studies expanding beyond short-read taxonomic profiling are relatively limited (30, 37).

The principal objectives of our study were to evaluate inoculation-based factors that may affect the establishment of gut bacteria in fruit flies and to evaluate the utility of near-full length 16S rRNA sequencing to track specific isolates. Specifically, our goals were to evaluate how adult fly age and diet inoculation formulation influence the colonization success of a target bacterial *Enterobacter* strain in medflies. This strain was originally collected from wild medfly larvae and transformed to exhibit antibiotic resistance and the florescent protein to provide ease of tracking this specific isolate. We evaluated inoculation success of this strain in both longstanding laboratory and mass-reared colonies of medfly. To complete our objectives, we used antibiotic resistance markers associated with the target strain to determine colonization success using plate counts on selective media with Kinnex full-length 16S rRNA PacBio sequencing of specimens to track our target strains and to evaluate the microbial diversity in the system.

## Methods

### Medfly Sources

Two rearing lines of medfly were used for these experiments, one considered laboratory and the other mass-reared. The laboratory colony is maintained at the USDA ARS at Daniel K. Inouye Pacific Basin Agricultural Research Center (DKI-PBARC) in Hilo, HI, USA. This colony (henceforth referred to as ‘PBARC’ colony) has been in culture for ∼20 years and fed on an artificial diet media following standard operating procedures (34). The mass-reared source of medfly (VIENNA 7 *tsl* strain) were obtained from the California Department of Food and Agriculture (CDFA) Fruit Fly Rearing Facility in Waimanalo, HI, USA. Medflies from this colony will henceforth be referred to as ‘CDFA’ flies.

### Bacterial Strain

Inoculations of flies were conducted using *Enterobacter sp.* CC2903LS21 originally isolated from field-collected medfly larvae obtained from infested coffee fruits. This wild-type strain was transformed with the florescent protein mScarlet and antibiotic selection markers tetracycline (Tet) and chloramphenicol (Chl) using established conjugation protocols using Tn5 transposon delivery system implemented in *E.coli S*17-1 (38, 39). pMRE-Tn5-165 was a gift from Dr. Mitja Remus-Emsermann (Addgene plasmid #118545; http://n2t.net/addgene:118545; RRID:Addgene 118545). Compared to wild-type cultures, this transformation also gives the colonies a distinct pink hue in addition to the antibiotic resistance.

### Age and Diet Formulation Evaluation Methods

The morning of the inoculation the target *Enterobacter* CC2903LS21 was grown at 32℃ in 2× lysogeny broth (LB) broth on a shaker until reaching an optical density OD600 of 0.5-1.0 (∼6h of growth time). The amount of CFUs present in the culture were estimated by using the OD reading coupled with the previously determined relationship between OD reading and CFU count of the target strain. Cultures were diluted in phosphate buffered saline (PBS, pH 7) to achieve a starting inoculum density of 10^6^ CFUs per mL or gram of diet, when added to the two diet formulations, described below.

To evaluate the impact of adult fly age on inoculation success, we compared flies that were new emerged adults (< 12h old) at time of inoculation to flies that were allowed to feed on the normal adult fly diet (a dry mixture of a 3:1 ratio of sucrose and yeast hydrolysate) for 7 days prior to inoculation.

To evaluate the impact of diet formulation on inoculation success we evaluated two different adult fly diet compositions referred to as the liquid diet and the slurry diet.

Nutritionally, the two diets contained the same ratio of sucrose and yeast hydrolysate, but the liquid diet was more dilute. Liquid diets were made using a 3:1 ratio of sucrose and yeast hydrolysate diluted and autoclaved in tap water (180 g L^-1^) and was autoclaved before use. The control flies received just this liquid diet. Flies that were inoculated target isolate received this liquid containing a dose of 10^6^ CFUs mL^-1^ of *Enterobacter* CC2903LS21. The liquid diet was provided to the flies by filling a microcentrifuge tube with hole bored close to the top with the liquid diet. A cotton wick placed under the bored hole provided a place where the flies could consume diet without entering the tube itself. The microcentrifuge tube was placed through a hole in the lid of the inoculation container (a plastic 450 mL round deli cup). A laboratory wipe was placed in the bottom of each container. The slurry diet formulation was made by mixing the normal adult fly diet (a dry mixture of a 3:1 ratio of sucrose and yeast hydrolysate) with a relatively small amount of liquid at a rate of 100 μL per gram of diet. This liquid to dry mixture ratio created a peanut butter like consistency. The inoculated flies received a dose of 10^6^ CFUs g^-1^of *Enterobacter* CC2903LS21 though this diet. Inoculations with this treatment were also done in plastic 450 mL round deli cups, however the cups had screened lids, and the flies were provided water through a small cup with a wick, in addition to the slurry diet.

PBARC flies were inoculated in mixed sex groups of 25-30 flies and replicated 3 times. CDFA inoculated in groups of containers of 15 and replicated two times. After the flies were set up in their inoculation containers, they were kept in a growth chamber set at 30℃ and 60-70% relative humidity. Flies were allowed to feed for 3 days, after which the flies were removed from the containers and processed. Flies were processed in one of two ways; 1) to determine the total and target strain CFUs counts, or 2) processed for full-length 16S rRNA sequencing, both processes are described in detail below.

Both the PBARC and CDFA flies were tested to see if there was difference between newly emerged and 7-day old flies, as well as between slurry and liquid diet. In addition, the CDFA flies had one more treatment group, flies that had been irradiated and flies that had not been irradiated. Flies that are used in the sterile male release program are irradiated before release, irradiation is done with gamma irradiation (150 Gy) by USDA APHIS located in Waimanalo. For the PBARC flies, both male and female flies were tested, whereas only males were tested with the CDFA flies. After the results with the PBARC flies, we tested newly emerged CDFA flies with both the slurry and liquid diet, however we only tested the 7-day old CDFA flies using the liquid diets.

### Retention of Colonized Bacteria Over Time

A supplemental study was done with newly emerged PBARC flies to evaluate the retention of *Enterobacter* CC2903LS21 within the fly after the initial inoculation period. Three time points after inoculation were evaluated: time zero (taken immediately after the inoculation period), one week, and two weeks. Flies were provided *Enterobacter* CC2903LS21 using the liquid diet as described above for three days. Flies for time zero were processed directly after the end of the initial inoculation period, whereas the rest of the flies were moved into their own individual cups to be held for the next one or two-week time period. These individual fly cups were small plastic cups (60mL) with a screened lid. A microcentrifuge tube with a hole and cotton wick was filled with water and given to the flies, as well as a small cup of dry 3:1 ratio sugar to yeast hydrolysate. The water tube was refreshed after one week for the flies that were being held for two weeks. At the end of their treatment period flies were processed to determine the total and target CFUs counts per fly were enumerated.

### Target Colony and Total Microbe Enumeration

For CFU enumeration, flies were surface sterilized submerging them in 10% bleach solution (Chlorox bleach) for 30 seconds followed by two 30 second rinses of autoclaved, distilled water. After surface sterilization, the flies were homogenized in 1ml of PBS. Fly homogenates were diluted and plated onto both LB agar (total colony counts) and onto LB agar containing 25 mg L^-1^ Tet and 15 mg L^-1^ Chl (target colony counts). This was done using an Eddy Jet Spiral Plater (Neutec Group Inc, Farmingdale, NY, USA). Plates were incubated for at least 16h before enumerating with a ProtoCol3 colony counter with software (Synbiosis, Frederick, MD, USA). Values were expressed as CFUs mL^-1^ fly^-1^. Some yeasts grew on the LB agar plates containing antibiotics, however the yeasts grew slower and had distinct morphological differences from the target bacterial stain, so they were able to be excluded from our counts.

### Full-Length 16S rRNA Sequencing

For 16S rRNA analysis, flies were surfaced sterilized identically to those for CFU determination, then homogenized in RNA/DNA Shield (Zymo Research, Irvine, CA, USA) and frozen at -80℃ until DNA extraction. DNA was extracted from samples using a Zymobiomics Magbead Kit (Zymo Research). Amplification of full-length 16S SSU rRNA was performed according to Pacific Biosciences (Melno Park, CA, USA) Kinnex 16S protocols with some modifications. Combinatorial indexed reactions were performed in 30 µL volumes consisting of 0.25 µM of the degenerative primers 27F (CTACACGACGCTCTTCCGATCT /10-mer barcode/ AGRGTTYGATYMTGGCTCAG) and 1492R (AAGCAGTGGTATCAACGCAGAG /10-mer barcode/ RGYTACCTTGTTACGACTT).

16S amplification was performed using NEB Q5 Hi-Fidelity Hot Start Polymerase (New England Biolabs) with the following conditions: 98℃ for 30s; 22 cycles of 98℃ for 10s, 55℃ for 15s, 72℃ for 2 min; and final extension of 72℃ for 10 min. Amplicons were pooled and magnetic bead purified prior to Kinnex library construction. Concatenated amplicons were produced according to the Kinnex 16S rRNA kit (Pacific Biosciences) and were sequenced on a Revio 24M SMRT cell on a Pacific Biosciences Revio sequencer. Read segmentation and demultiplexing of sequences were conducted using SMRTLink v.13 software (Pacific Biosciences).

Un-segmented and demultiplexed HiFi reads were processed for downstream statistical analysis using a nextflow pipeline designed for full-length 16S data (pb-16S-nf v 0.7; https://github.com/PacificBiosciences/HiFi-16S-workflow) on USDA ARS SCINet HPC Resources. Sequences were pre-screened for quality scores >Q20 and primers were trimmed from reads (40, 41). Data were processed with dada2 implemented in QIIME 2 using default parameters except using -min_length 1300 (42, 43). Denoised ASV sequences were classified outside of the pipelines parameters using the RDP Naïve Bayesian Classifier with using RDP Database v19 (44).

After our main analysis, we compared full-length 16S rRNA with the V4 and V3V4 subunits by extracting them *in silico*. Samples were processed as described above, and after quality filtering, the expected subregion was extracted with primers 515F and 806R for V4 (45, 46) and 341F and 806R for V3-V4 (47) in dada2. Following initial extraction, all subsequent steps were identical to those implemented in the full-length pipeline, with the only changes being the expected length of the fragments during the filtering steps.

### Data Analysis

Statistical analyses were performed in R Studio (48) using R version 4.4.1 (49). Explanatory variables were different between our specific experiments but were all analyzed using a similar strategy using non-parametric tests. All CFU data were analyzed using pairwise Wilcoxon rank sum tests using Holm-corrected p-values.

16S rRNA ASVs were subsampled to a value to ensure that >90% of the samples receiving reads were included in the analysis to account for bias in sequencing depth. This number did vary between the specific tests. Comparisons between PBARC colony reared flies and CDFA flies were performed with data subsampled to 15,000 reads. Rarefaction was performed using the R package vegan v.2.6-8 (50). After subsampling, we computed Bray-Curtis dissimilarities between samples in vegan. Non-metric dimensional scaling was performed for each experiment. Permutational multivariate analysis of variance (PERMANOVA) and pairwise PERMANOVA (51) were performed separately for each experiment. For our experiments evaluating bacterial establishment with the PBARC colony, we included differences with sex, fly age, and diet formulation as fixed effects in our overall model. We did not observe overall differences in sex, so we pooled those groups together for pairwise comparisons between groups. Similarly, the mass-reared CDFA colony, we initially included irradiation, diet formulation, and inoculation in our initial analysis, but irradiation did not have an overall statistical effect or exhibit interactions with other variables so was pooled for purposes of pairwise comparisons.

Sequencing of *Enterobacter* CC2903LS21 alongside our samples yielded six dominant ASVs (Supplemental Table 1). We computed the total relative abundance of the *Enterobacter* CC2903LS21 signature ASVs for each sample and performed Wilcoxon rank sum tests. To further evaluate relationships between *Enterobacter* in our experiments, we constructed stacked barcharts of the relative abundances of ASVs grouped at the genus level. Additionally, we constructed heat maps of ASVs classified as *Enterobacter* using z-transformed relative abundances of individual ASVs with the R package pheatmap v.1.0.12 (52). Clustering of distances on heatmaps were conducted using Bray-Curtis dissimilarities between samples and Manhattan distances between ASVs using complete linkages.

The following packages also were used to facilitate our analyses at various steps: ggpubr 0.6.0 (53), ggh4x v.0.7.5 (54), ggtext v.0.1.2 (55), goeveg v.0.7.5 (56), rstatix v.0.7.2 (57), and tidyverse v.2.0.0 (58).

## Results

### Near Full-Length 16S rRNA Sequencing Metrics

PacBio Kinnex technology boasts increased sequence yields compared to the existing SMRT-bell adapter-based methods by concatenating twelve amplicons into final library molecules approximately 17.5 kb in length, increasing the output 12-fold. After sequencing the chimeric library is desegmented into the amplicon subunits. Our total run yielded 28.7 million desegmented, demultiplexed circular consensus sequence (HiFi) reads with 95% of those reads were >Q20 quality. After primer trimming, 26.8 million remained (93% of HiFi reads) for filtering in dada2. There were 20.1 million denoised reads where of those denoised of which they were 96% non-chimeric. Through all filtering steps, 67% of the 28.7 million read input yielded high quality, filtered, non-chimeric ASVs. For the samples we analyzed here, we obtained 14 million reads for the experiments. There was a range in per sample sequencing depth (<1,000 to ∼500,000) that was indicative of non-uniform pooling efficiencies. Samples with sequences lower than 15,000 were excluded from subsequent analyses.

### Impact of Diet and Age on *Enterobacter* Establishment and Microbiome of PBARC Flies

Medfly age coupled with diet formulation affected the establishment of *Enterobacter* in adult flies (Figure 1). Fly inoculations were generally successful in liquid formulations, with Tet and Chl resistant colonies being observed in both newly emerged and flies that were seven days old (Figure 1). Contrary to our initial expectations, we saw no differences in the establishment success between age classes for the liquid diet (*W* = 434; n=30; p = 0.813). However, for the slurry inoculation, we observed significantly less establishment in the older flies compared to newly emerged flies (*W* = 754; n_1_=29 n_2_=30; p <0.001). Total counts exhibited comparable trends, where differences were not present in liquid formulation (*W* = 555; p = 0.122), but seven-day old flies had slightly lower titers (∼10%) compared to younger for the slurry method of inoculation (*W =* 644, p = 0.002). Comparing across diet formulations, liquid and slurry did not differ for newly emerged flies (*W* = 527; n=30; p = 0.167). We did not observe differences between male and female flies on the titers under any of the conditions (p > 0.05). In assessing non-inoculated controls, we had two incidences of colonies growing on antibiotic plates (Supplemental Figure 2), and both were at levels below the threshold for establishment. Some newly emerged colonies exhibited extremely low titers (< 200 CFUs) on LB plates for both slurry and liquid diets. Older flies had titers between 10^7^ – 10^8^ per fly, which is typical for these lines (Supplemental Figure 1) and matches our inoculation results (Figure 1).

**Figure 1:**
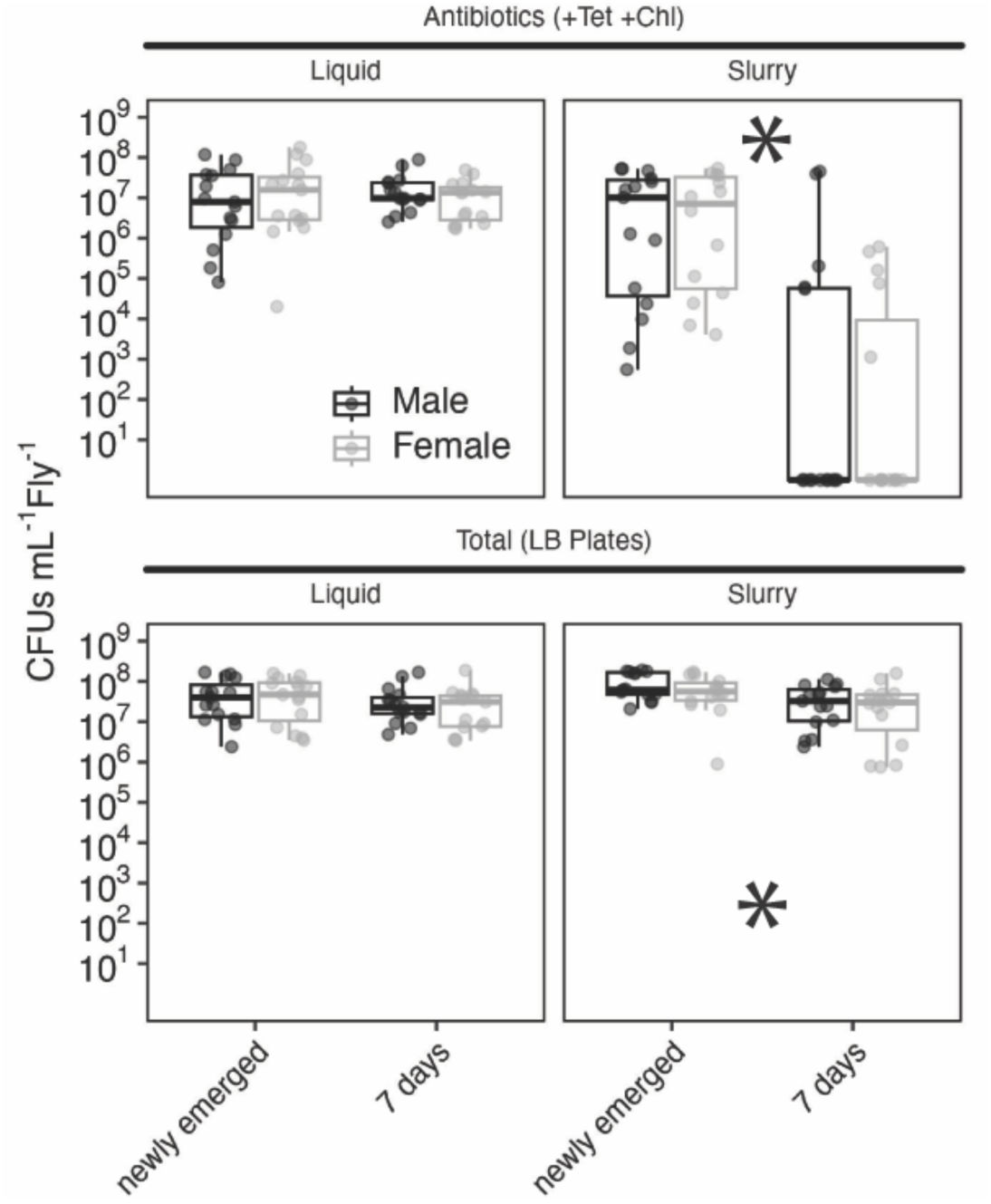
Colony forming units (CFUs mL^-1^ fly^-1^) of USDA ARS reared medfly by *Enterobacter* CC2903LS21-Tn165 resistant to chloramphenicol and tetracycline. Newly emerged flies or flies that were seven days old were inoculated using two different diet formulations, liquid or slurry. Each circle represents colony counts from an individual. Asterisks denote significant differences (p < 0.05) between groups. Experiments were performed across three cohorts. No differences in establishment were observed between males and females.

Differences between groups were further evaluated using near full-length 16S rRNA sequencing (Figure 2). Non-metric dimensional scaling of Bray-Curtis distances exhibited clear separation of sample types in two-dimensional space (Figure 2A). PERMANOVA indicated strong and interactive effects of diet (R^2^ =0.14; F=81.4; p <0.001), inoculation (R^2^ =0.26; F=153.9; p <0.001), and age (R^2^ =0.22; F=131.0; p <0.001) on the ASV composition (all interaction had p <0.001; full statistics with interactions in Supplemental Table 1). Pairwise comparisons indicated separation between each sample group (Supplemental Table 2), with the exception being between seven-day-old slurry control and slurry *Enterobacter* (Figure 2A).

**Figure 2:**
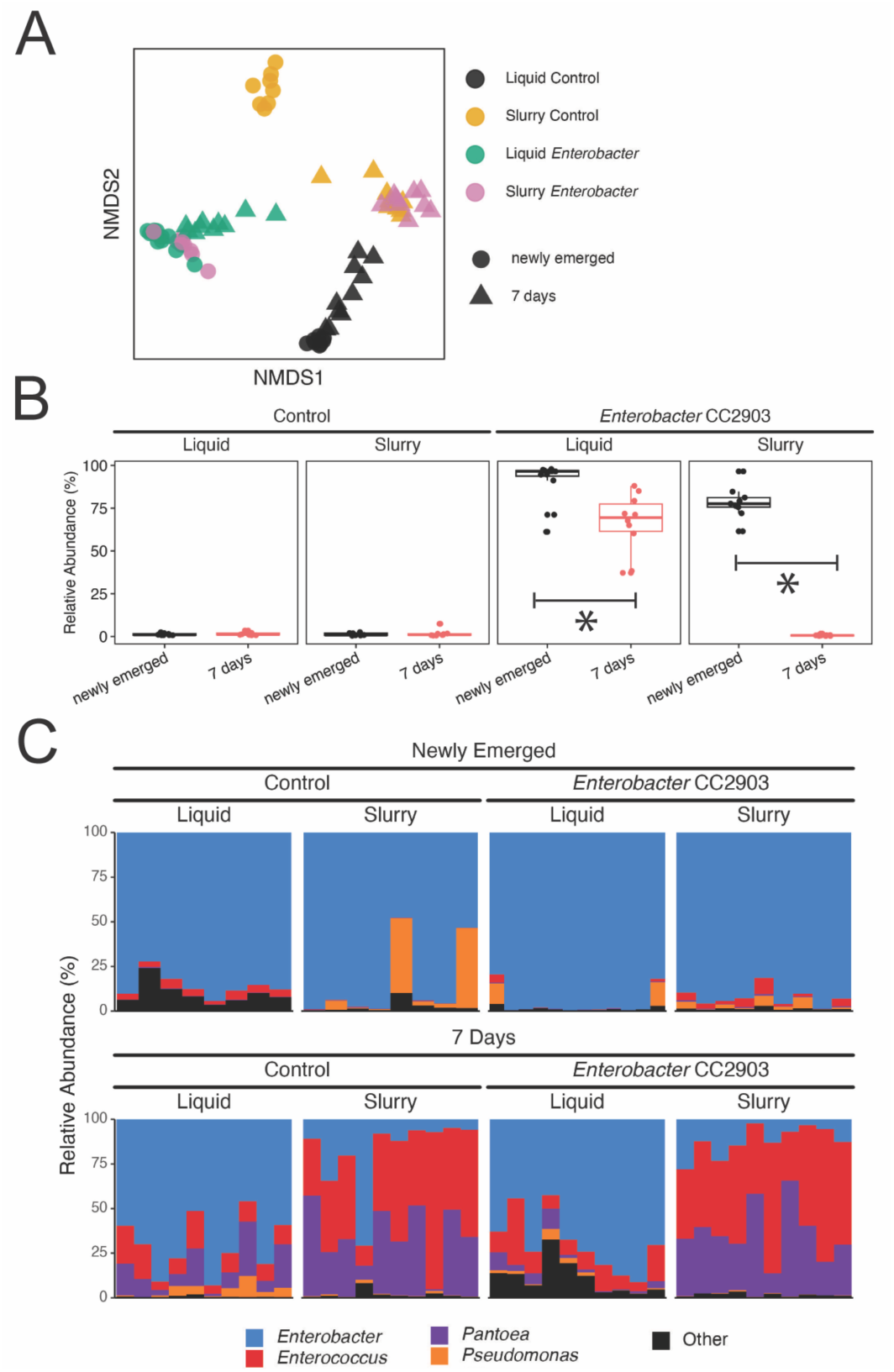
Full-length 16S-rRNA sequencing of control and *Enterobacter* CC2903-inoculated PBARC medfly at two age classes and two diet formulations. (A) Non-metric multidimensional scaling plot (NMDS 2D Stress = 0.12) of control and inoculated samples. Colors denote diet- inoculation combination and shape represents age class. Pairwise PERMANOVA differences between samples are reported in Supplemental Table 1. (B) Relative abundance of the five ASVs correspond to *Enterobacter* CC2903 in control and inoculated samples. Asterisks denote significant differences between samples (p < 0.05). (C) Stacked bar charts of the relative abundance of taxonomic classifications at the genus level. ‘Other’ includes taxa at the genus level comprising < 2% of the read relative abundance across all samples.

Relative abundance of the *Enterobacter* CC2903LS21 ASVs differed between samples (Figure 2B). In flies inoculated with *Enterobacter*, there were significantly greater relative abundances of the target *Enterobacter* in newly emerged compared to seven-day-old for both liquid (*W* = 108, n_1_=12 n_2_=10; p <0.001) and for slurry diets (*W* = 90, n_1_=9 n_2_=10; p <0.001). Newly emerged flies inoculated with liquid diet had higher relative abundances than those inoculated with slurry diet (*W* = 87, n_1_=12 n_2_=9; p <0.018).

Stacked barplots of genus level taxonomy demonstrated some interesting trends (Figure 2C). Newly emerged flies, regardless of being inoculated with the target *Enterobacter* CC2903LS21 or not, were dominated by ASVs classified as *Enterobacter* by the RDP database. Further, both control and inoculated seven-day-old flies fed liquid had high incidences of *Enterobacter* classification (< 75%). Older flies fed slurry, either with or without *Enterobacter* CC2903LS21, exhibited greater diversity. Those older flies had elevated relative abundances of both *Enterococcus* and *Pantoea* relative to the other samples, with *Pseudomonas*, *Enterobacter*, and less abundant taxa rounding out the composition.

### Impact of Formulation on *Enterobacter* Inoculation in Newly Emerged Mass-Reared Flies

*Enterobacter* inoculation of newly emerged mass-reared CDFA medfly differed between diet formulations (Figure 3A). Resistant colonies were in much higher incidence and titer in flies inoculated with liquid diet compared to slurry diet (Figure 4A; *W* = 325, n_1_=21 n_2_=16; p<0.001). No differences were observed in total plate counts (*W* = 114, n_1_=21 n_2_=16; p =0.098). For control processing (Supplemental Figure 3), we observed some instances of colony growth (1/12 10^5^; 3/12 ∼10^2^). Visual inspection suggested they were potential yeasts. All these values were below the inoculation rate of the controls, and did not have the morphology or color or the target colony. Total counts in control inoculations ranged from 10^7^ – 10^8^ per fly.

**Figure 3:**
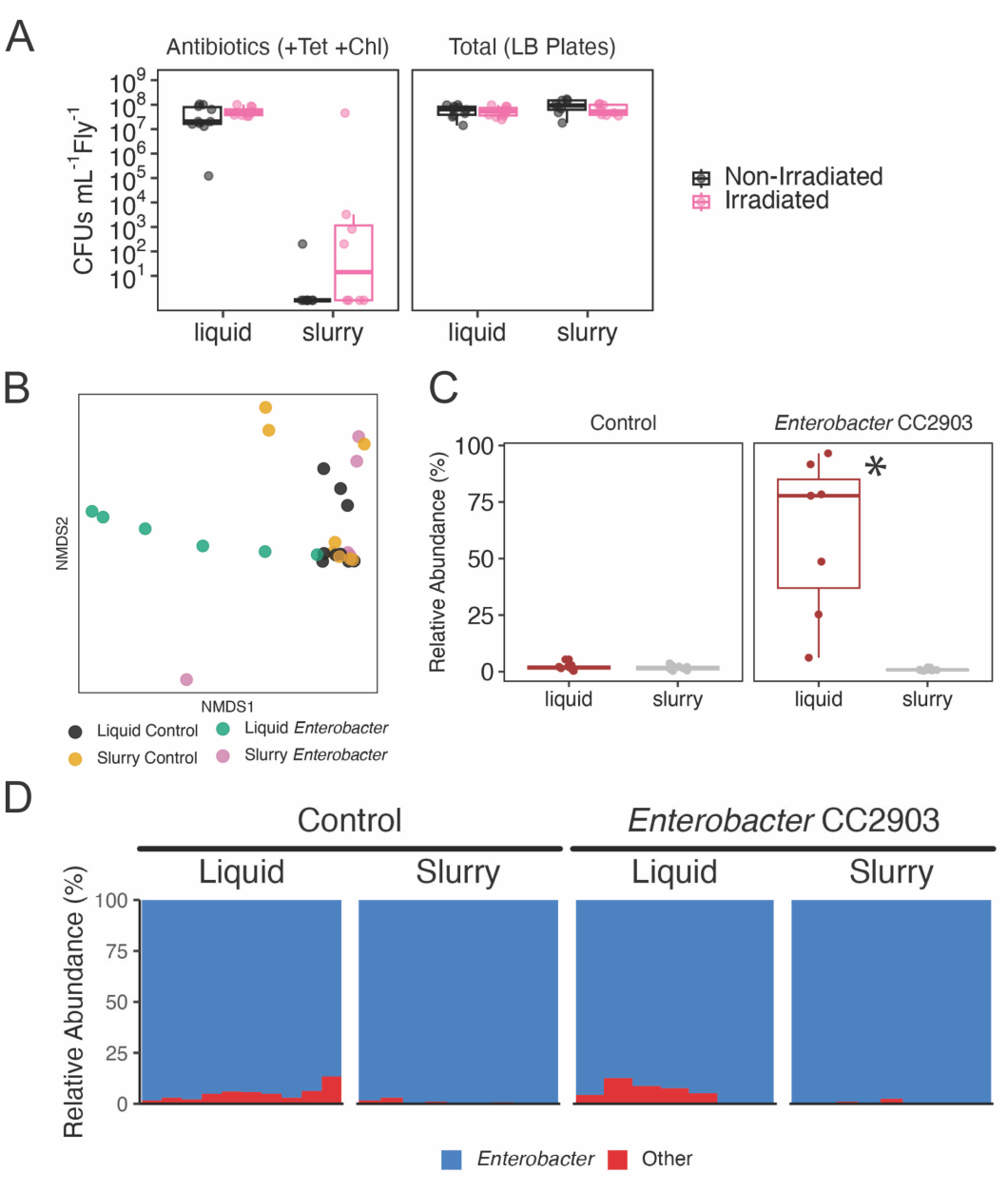
Influence of diet formulation on bacterial establishment in mass-reared fruit flies. (A) Colony forming units (CFUs) mL^-1^ fly^-1^ of mass-reared medfly inoculated with *Enterobacter* CC2903LS21-Tn165 resistant to chloramphenicol and tetracycline. No differences in establishment were observed between irradiated and non-irradiated flies based on CFUs. (B) 16S-rRNA sequencing of control and *Enterobacter* CC2903-inoculated CDFA medfly at two diet formulations with non-metric multidimensional scaling plot (NMDS 2D Stress = 0.14) of control and inoculated samples. Colors denote diet-inoculation combinations. Pairwise PERMANOVA differences between samples are reported in Supplemental Table 2. (C) Relative abundance of the five ASVs correspond to *Enterobacter* CC2903 in control and inoculated samples. Asterisks denote significant differences between samples (p < 0.05). (D) Stacked bar charts of the relative abundance of taxonomic classifications at the genus level. ‘Other’ includes taxa at the genus level comprising < 2% of the read relative abundance across all samples.

**Figure 4:**
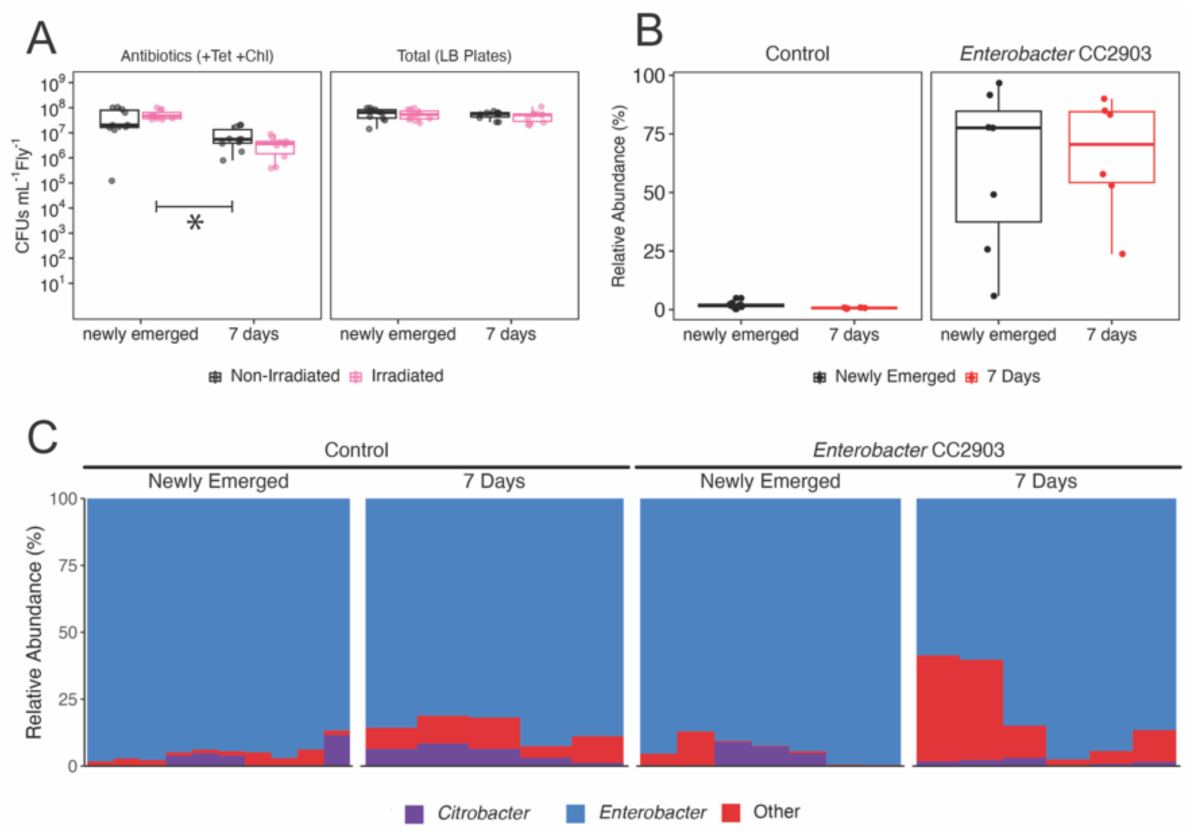
Influence of age on establishment of bacteria using a liquid diet formulation in mass- reared medfly. (A) Colony forming units (CFUs) mL^-1^ fly^-1^ of mass-reared medfly inoculated with liquid diet with *Enterobacter* CC2903LS21-Tn165 resistant to chloramphenicol and tetracycline at two different ages. (B) 16S-rRNA relative abundance of the five ASVs correspond to *Enterobacter* CC2903 in control and inoculated samples. Asterisks denote significant differences between samples (p < 0.05). (C) Stacked bar charts of the relative abundance of taxonomic classifications at the genus level. ‘Other’ includes taxa at the genus level comprising < 2% of the read relative abundance across all samples.

Full-length 16S rRNA sequencing supported differences in inoculation procedures (Figure 3B). We observed separation of the liquid *Enterobacter* inoculation in two-dimensional space (Figure 3B). PERMANOVA indicated significant effects of diet formulation (R^2^ =0.098; F=6.52; p <0.001), inoculation (R^2^ =0.16; F=10.9; p <0.001), and their interaction (R^2^ =0.11; F=7.15; p <0.001) on ASV composition. Pairwise comparisons (Supplemental Table 3) demonstrated significant differences between the liquid *Enterobacter* treatment from all others (R^2^ > 0.38; p <0.001), while no other differences were observed.

ASVs corresponding to *Enterobacter* CC2903LS21 were significantly higher in liquid-fed flies than in slurry-fed (Figure 3C; *W* = 96; n_1_=7, n_2_=9; p= 0.002). However, taxonomic profiling indicated a high relative abundance of sequences classified as *Enterobacter* across all samples (Figure 3D), like in PBARC-reared flies.

Inoculation was successful in older CDFA flies using liquid (Figure 4A). Target *Enterobacter* cultures were established in both newly emerged and in older flies, but older fly titers were reduced by ∼10% (*W* = 388; n_1_=21, n_2_=20; p<0.001). No differences in total culturable titers were observed between the age classes (*W* = 247; n_1_=21, n_2_=20; p= 0.341).

However, 16S rRNA sequencing indicated no differences were seen in the target *Enterobacter* CC2903 ASVs (Figure 4B; *W* = 20; n_1_=7, n_2_=6; p= 0.945). Like with the other sequencing datasets, regardless of inoculated with a target *Enterobacter*, most of the sequences were classified as *Enterobacter* with the RDP database (Figure 4C).

### Full-Length 16S rRNA Sequences Showcases *Enterobacter* Diversity in Medfly Colonies

Comparing sequences classified as *Enterobacter* by RDP of newly emerged flies from our experiments indicated ASV distributions and suggested different strains associated with the different fly sources (Figure 5). Besides the inoculated strain (I), there were seven additional clusters of abundant ASVs distributed across the samples. These clusters were distributed across the different sample types. There were three clusters completely associated with PBARC flies (II, III, IV) and two major cluster that were associated with CDFA flies (V, VIII). Less frequent clusters were associated with one or two individual flies (VI, VII).

**Figure 5:**
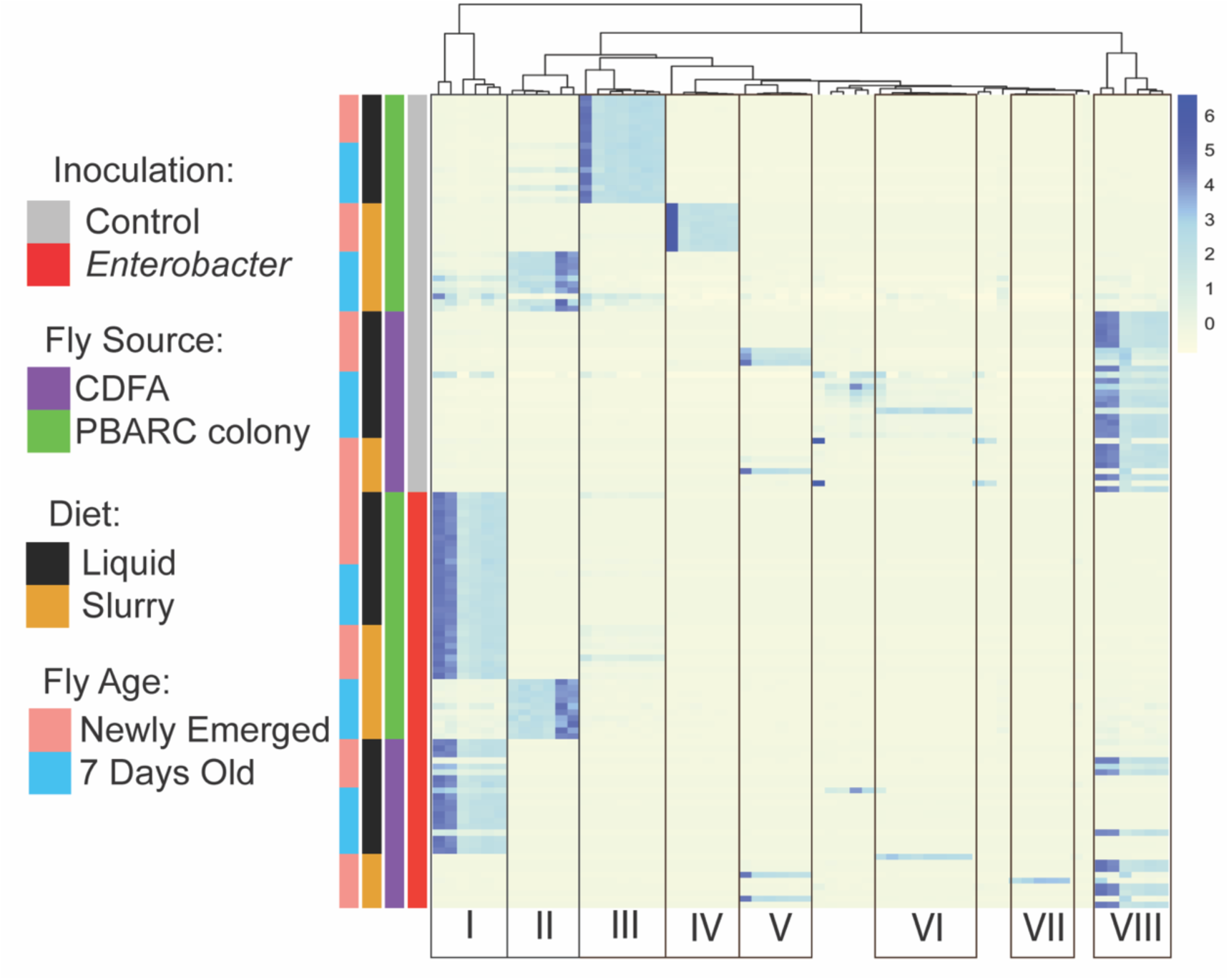
Relative abundances (z-transformed) of individual ASVs classified as *Enterobacter* at the genus level via the RDP database across all samples for experiments with CDFA and PBARC medfly colonies. Clustering of ASVs was conducted using complete method of Manhattan distances. Samples (rows) are not clustered based on distances. Roman numerals denote main clustered ASVs (Cluster I = ASVs comprising *Enterobacter* CC2903LS21).

When we evaluated shorter fragments by extracting V4 (Supplemental Figure 4) and V3- V4 (Supplemental Figure 5), much of this signal was reduced. For V4, the Enterobacteriaceae sequences collapsed into a single ASV (Supplemental Figure 4C). V3-V4 provided more ASV diversity, but did not have clear patterns like with full-length heatmap (Supplemental Figure 5C).

### Target Isolate Retention in Medfly Following Inoculation

Target bacterial isolates remained established in individually reared medfly following inoculation with *Enterobacter* CC2903LS21 (Supplemental Figure 6). Resistant bacteria were consistently detected immediately following inoculation (time zero), one week, and two weeks following treatment at levels exceeding 10^6^ CFUs mL^-1^ per fly. We observed decreases in antibiotic resistant colonies between time zero and one week (*W* =94; n=10; p. <0.001) but saw no differences between the other timepoints (p > 0.1). Total colony counts followed similar trends, where one week was significantly lower (p < 0.04) compared to time zero and two weeks after inoculation.

## Discussion

Reintroduction of field-collected microorganisms to mass-reared fruit flies has a potential to improve the performance of males reared for SIT (20, 21, 59). Here, our principal objectives of this study were to determine how fly age and inoculation formulation through diets influence the establishment of target microbial associates. Using an *Enterobacter* strain isolated from wild medfly stably transformed with two antibiotic resistance markers, we determined that we could introduce this isolate to independent medfly rearing lines using a liquid formulation. Full-length 16S rRNA sequencing corroborated these results, especially as we were able to track a multi- ASV signature unique to our introduced *Enterobacter*. This sequencing uncovered an initially unexpected observation related to sequences classified as *Enterobacter*; groups of *Enterobacter* ASVs formed distinct clusters across sample types suggesting potentially unique strains and greater diversity than initially meets the eye even among laboratory reared populations.

Our overarching hypothesis was that fly age would impact bacterial establishment in the gut. This was grounded in the fact that we see significant differences in microbial titers as medfly adults age (34), similar to other insect species following adult emergence (60). We expected individuals possessing higher bacterial titers would be more resistant to colonization than those with lower titers due to competitive exclusion (9, 10, 61). Our slurry diet formulation had this anticipated result, with there being little to no colonization of our target *Enterobacter* in flies that with gut bacterial densities exceeding 10^4^ CFUs mL^-1^ fly^-1^. Lack of colonization may not solely due to bacterial titers, as there may also be age-dependent changes in the gut defenses (62, 63) and differences in diet ingestion rate that may also be at play in mediating bacterial colonization and establishment.

Our results with liquid inoculum yielded contrasting results to the slurry diet. Age had little impact on how well the bacteria established when it was introduced by a liquid formulation. This may be related to bacterial propagation in the liquid, and the density that was ingested by the flies. Measuring fly consumption of food is challenging in our experimental context, but we presume there may be more liquid consumption than solid consumption due to concentrations of nutrients. This can alter feeding behaviors. For example, in the Queensland fruit fly, *Bactrocera tyroni*, dilution of adult diet with water resulted in the ingestion of greater volumes of diet (64), likely a form of compensatory feeding to obtain its required intake of nutrients (65). Even if a microbial community is initially resilient to bacterial introduction, high enough inoculum density may overtake an established community (66). While we controlled our initial inoculum density, we did not account for fluctuations over time within the liquid media to keep everything constant. So, while the diets may have initially similar colonization densities, the different formulations may differ in how many are supported. Our results with a liquid formulation support those of a prior study in medfly fed 10^9^ CFUs mL^-1^ streptomycin-resistant *Klebsiella oxytoca* liquid culture (23), where the introduction of bacteria was stable and detected for days following inoculation. Overall, these findings are not necessarily negative, since these results show a liquid inoculation is fly source/strain and host age agnostic. However, subsequent resilience and how the gut microbiome may change in the field warrants further inquiry. This may be a consideration to investigate moving forward to understand the minimal concentrations and the amount of ingestion required to override established gut microbiomes.

Most of the sequences were classified as *Enterobacter* with the RDP sequence database at the genus level, regardless of targeted treatment or fly source. We were able identify that *Enterobacter* CC2903LS21 has a high percentage of six ASVs (Supplemental Table 1).

Enterobacteriaceae often possess multiple 16S rRNA gene copies that may vary in total numbers between strains and in intragenomic sequence within a strain (67–69). Interestingly, we saw variable patterns of *Enterobacter* ASV association between individuals across different diet treatments and locations, suggesting different species (or strains) (70, 71) colonize and occupy these systems under different environmental conditions. This variation is unlikely to be explained by error or random chance, as typically only a single *Enterobacter* cluster was primarily associated with an individual (Figure 5). When we investigated ASV distribution using 16S rRNA sequences (V4 or V3V4; Supplemental Figures 4 and 5), the ASV partitioning and potential species or strain-level variation was totally obfuscated.

These results were initially unexpected, or at least not an initial focus of our study, but provide some important insight into potentially greater microbial diversity in insect systems. Even at the genus level, Enterobacteriaceae isolates can be notoriously difficult to parse with full-length 16S rRNA (72–75). Partial 16S rRNA sequences has been shown exacerbate this issue for some genera in this clade (76). This is noteworthy since several plant-feeding insects possess have robust interactions with commensal gut microbiota within the Enterobacteriaceae, including fruit flies (16), caterpillars (77, 78), and some beetle lineages (79–81).

Our data support the notion full-length 16S rRNA ASVs afford better tracking of isolate establishment, turnover, and maintenance in these insects as opposed to fragments. Historically, full-length 16S rRNA amplicon generation was more costly per read, but through implementation of Kinnex technology on a Revio platform, the cost was on par, or lower than 600 cycle-based Illumina approaches. Like other studies, our results indicate that the full-length approaches can at a minimum improve taxonomic classification resolution (82–86). Whether full- length 16S rRNA outperforms sub-regions (like V4 or V3-V4) for beta-diversity metrics may depend more upon within-sample composition alongside variation in the sequences (82, 84). Demonstrated here, communities with a dominant group belonging to the Enterobacteriaceae may pose problems with some 16S rRNA subregions. As discussed in detail by others (71), caution should still be taken using ASV approaches in terms of diversity-based metrics (87) and whether ASVs should be favored over more traditional operational taxonomic unit (OTU)-based approaches for these metrics (88). On one hand, ASVs splitting intragenomic variation in 16S rRNA can have inconstant effects on diversity metrics as copy number and intragenomic variation can vary between isolates and strains (87, 89, 90). Alternatively, splitting of ASVs combined with isolate-level resolution can help determine strain-specific dynamics in well-defined experiments (71). Overall, these technical trade-offs should be assessed on a case-by-case basis and informed by the overall study design.

Although our experiments provide guidance on some factors that influence the resiliency of the medfly microbiome, there are some caveats. In our initial design, our experiments only used one isolate to track, and future work should consider more strains. Our study employed two different fly sources, but we caution direct comparisons between the two. They are from different rearing conditions and have different genetic backgrounds. Controlling for these differences is a step that needs to be taken to understand how fly strain and different gut microbiome strains can influence the function and resiliency of the system. We also did not conduct any performance measures of the strains on our system but are certainly components to be addressed in the next stages of this research. Similarly, we have no basis to speculate on potential differences in function of the *Enterobacter* strains we have observed in our full-length 16S rRNA dataset.

Overall, our study provides some practical guidance alongside other avenues for further investigation. First, liquid diet being efficient to inoculate flies regardless of colony source or age is an important result. These observations indicate we do not necessarily need to be concerned with starting microbial titer to introduce different bacteria to adult medfly males consistently from mass-reared locations. Further, our inoculation procedures provided consistent results between sources, as well as between mass-reared male flies that were both irradiated and non-irradiated. We observed that our focal bacteria was present in flies reared individually for at least two weeks following introduction in a growth chamber (Supplemental Figure 6), supporting prior observations (23). Next steps would involve determining longevity and resiliency of inoculated flies under semi-natural conditions, such as in field cages with plants under ambient environmental conditions. Using an rDNA-expressing strain would not be an appropriate tool to implement outside of the laboratory from regulatory perspective, but long-read 16S rRNA sequencing suggests we can track specific wild-type Enterobacteriaceae strains using a multi-copy approach. Outside of antibiotic resistance (23), determination of successful inoculations of specific strains has not been widely assessed in tephritids (36). While this approach needs to be validated, this may be a good tool to facilitate an improved understanding of the dynamics and fluctuations of functionally important, but difficult to resolve Enterobacteriaceae with 16S rRNA that are prevalent among plant-feeding insect pests.

## Acknowledgements

We thank Jean Auth for assistance in the initial cultivation of the *Enterobacter* strain used in the study. The findings and conclusions in this publication are those of the authors and should not be construed to represent any official USDA or U.S. Government determination or policy. Mention of trade names or commercial products in this publication is solely for the purpose of providing specific information and does not imply recommendation or endorsement by the U.S. Department of Agriculture. USDA is an equal opportunity provider and employer. This research used resources provided by the SCINet project and/or the AI Center of Excellence of the USDA Agricultural Research Service, ARS project numbers 0201-88888-003-000D and 0201-88888- 002-000D. This research was supported in part by funding from USDA-APHIS by Plant Protection Act 7721 to CJM (58-2040-3-021-IA) and by the U.S. Department of Agriculture, Agricultural Research Service appropriated project “Advancing Molecular Pest Management, Diagnostics, and Eradication of Fruit Flies and Invasive Species” (2040-22430-028-000-D). IS was funded by the USDA National Institute of Food and Agriculture (NIFA) Hatch project (HAW09051-H), managed by the College of Tropical Agriculture and Human Resources, University of Hawaii at Manoa.

## Data Availability

Raw sequences have been submitted to NCBI SRA under the accession number PRJNA1173595.

