## Supplemental Files for "Utilizing full-length 16S rRNA sequencing to assess the impact of diet formulation and age on targeted gut microbiome colonization in laboratory and mass-reared Mediterranean fruit flies"

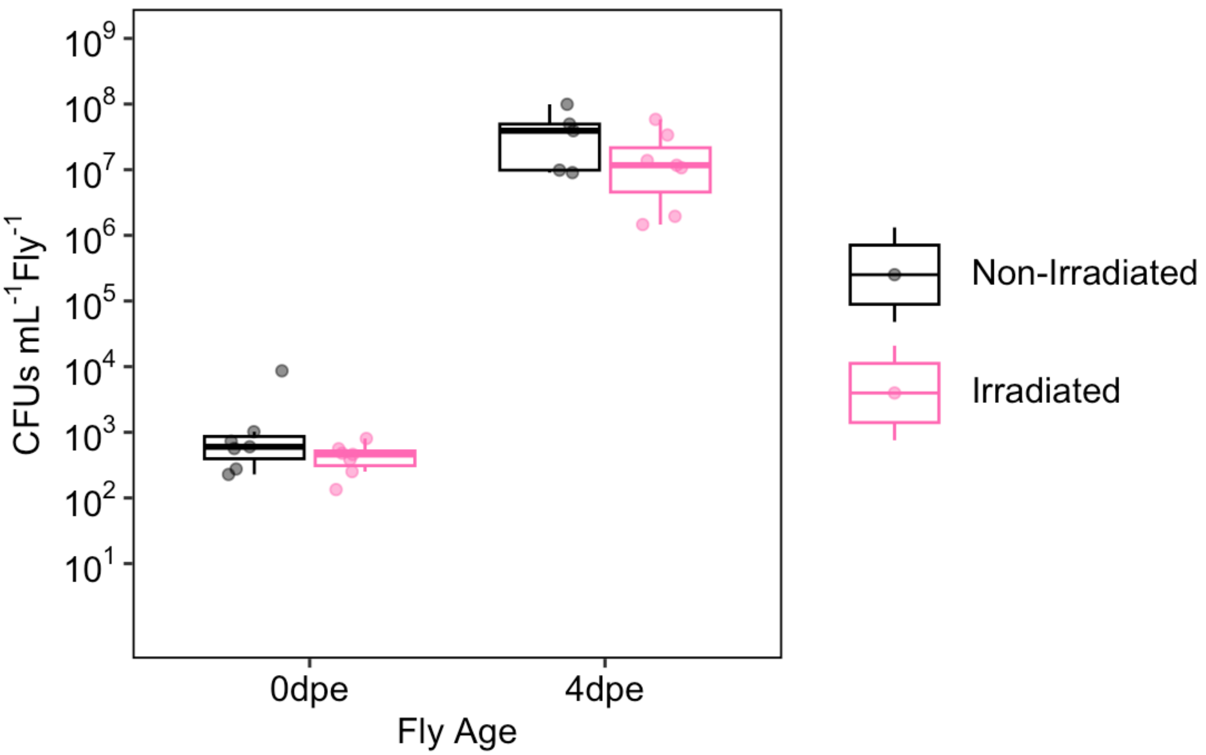


Supplemental Figure 1: Culturable (LB) titers of CDFA flies at adult emergence (0dpe) and four days post adult emergence (4dpe).

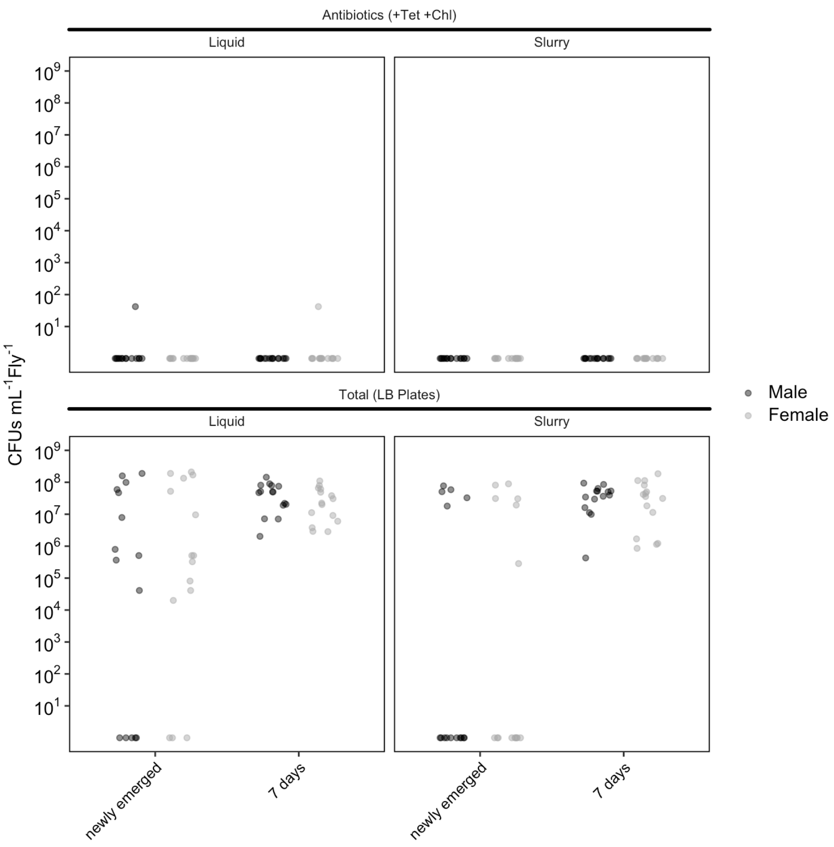


Supplemental Figure 2: Detection of bacteria from PBARC control inoculated flies. Flies were quantified three days after feeding on broth.

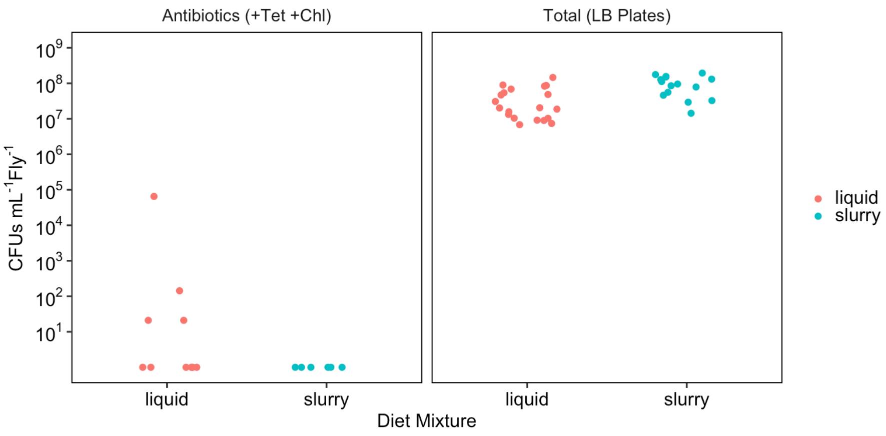


Supplemental Figure 2:
Growth of bacteria from CDFA control inoculated flies. Flies were quantified three days after feeding on broth.

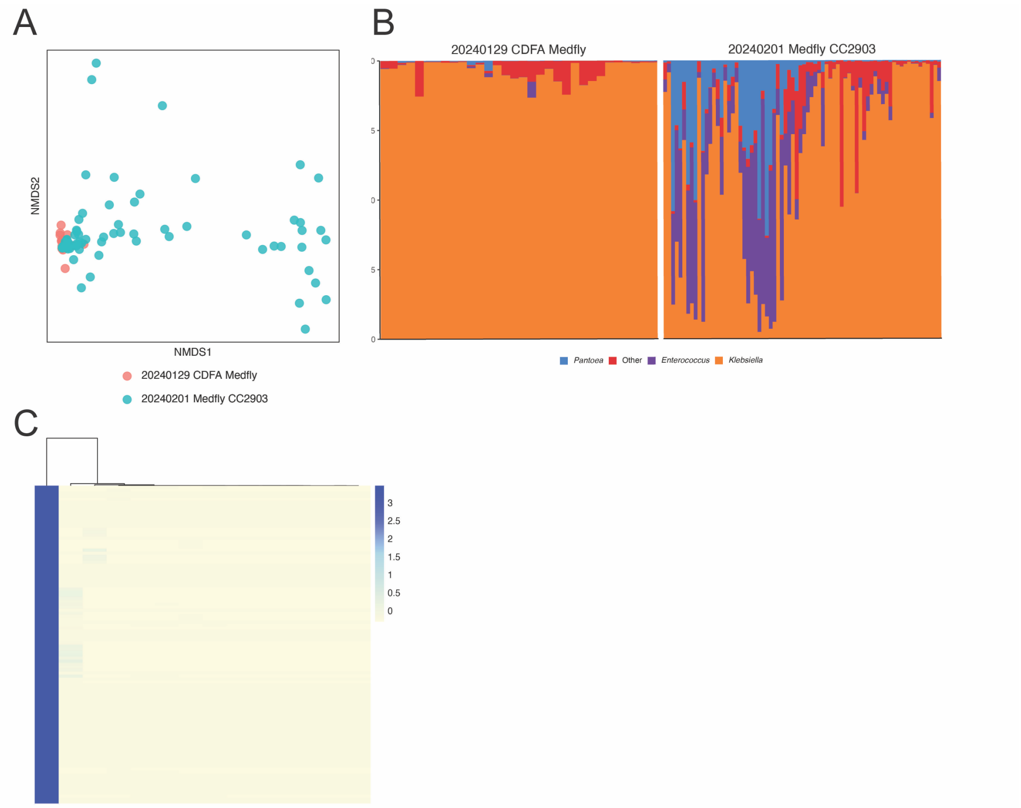


Supplemental Figure 4: Evaluation of community members with the V4 region of the 16S. V4 was extracted *in silico*. (A) NMDS of the two experimental blocks (CDFA and PBARC). (B) Stacked barcharts at the genus level. (C) Heatmap of ASVs classified at as Enterobacteriaceae.

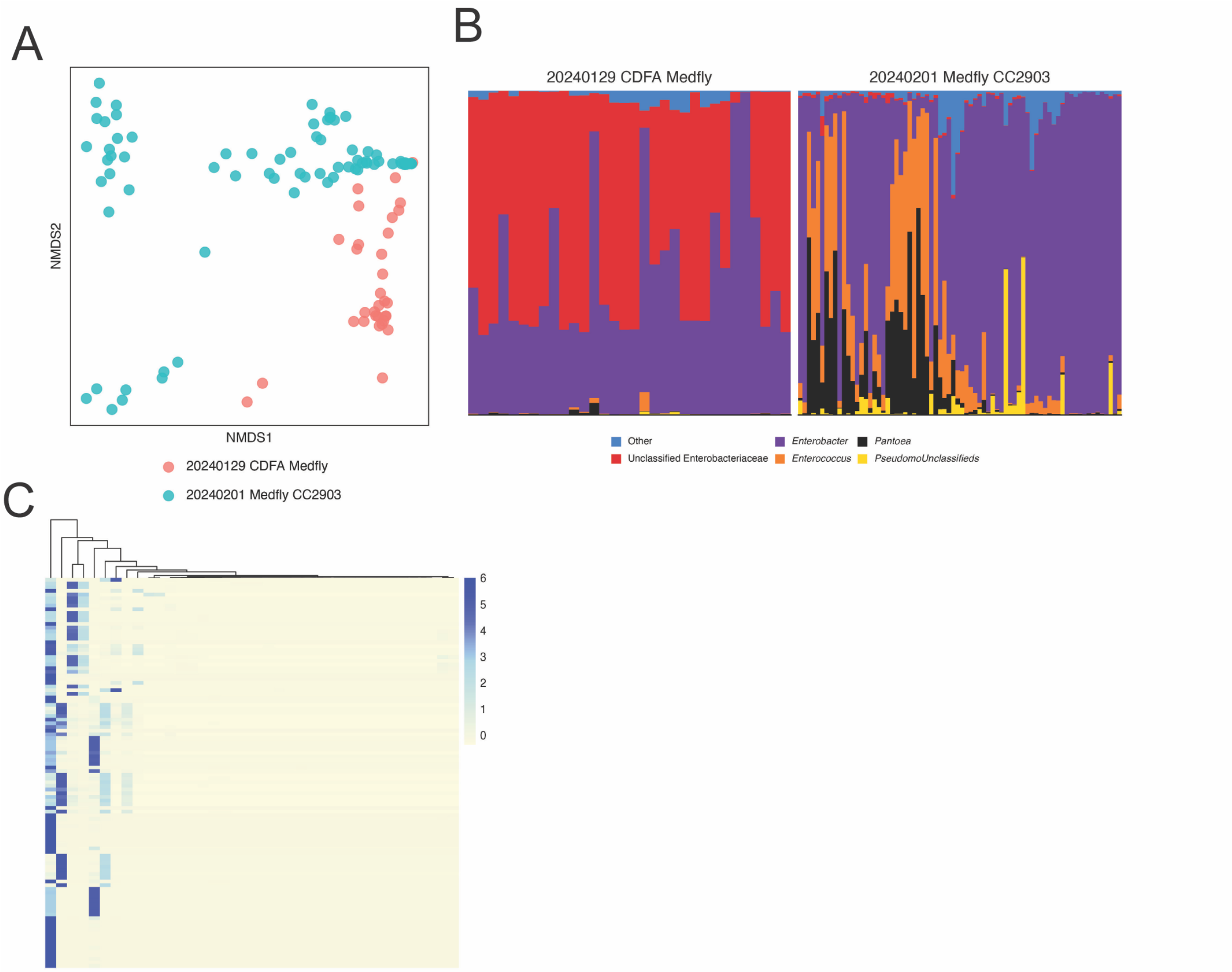


Supplemental Figure 5: Evaluation of community members with the V3-V4 region of the 16S. V3-V4 was extracted *in silico*. (A) NMDS of the two experimental blocks (CDFA and PBARC). (B) Stacked barcharts at the genus level. (C) Heatmap of ASVs classified at as Enterobacteriaceae.

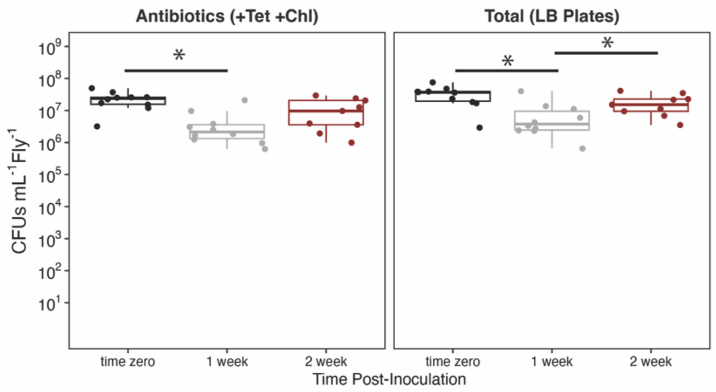


**Supplemental Figure 5**: Changes in detection of antibiotic-resistant colonies in flies inoculated with *Enterobacter* CC2903LS21. Asterisks represent significant differences between groups ($p < 0.05$).

Supplemental Table 1: ASV composition of pure culture of *Enterobacter* CC2903LS21

| | *Enterobacter* CC2903LS21 |
|---|---|
| Total Reads | 3859 |
| % ASV_1 | 24.95 |
| % ASV_2 | 24.75 |
| % ASV_5 | 13.63 |
| % ASV_6 | 11.66 |
| % ASV_7 | 11.64 |
| % ASV_10 | 12.52 |
| % Total | 99.20 |

Supplemental Table 2: Pairwise PERMANOVA on medfly microbiome for PBARC experiment

| Comparison being Tested | | $R^2$ | F | p-value |
|---|---|---|---|---|
| Between Newly Emerged: | Liquid Control vs Slurry Control | 0.891 | 113.8 | 0.002 |
| | Liquid Control vs Liquid CC2903 | 0.948 | 328.5 | <0.001 |
| | Liquid Control vs Slurry CC2903 | 0.943 | 250.1 | <0.001 |
| | Slurry Control vs Liquid CC2903 | 0.884 | 137.4 | <0.001 |
| | Slurry Control vs Slurry CC2903 | 0.879 | 109.6 | <0.001 |
| | Liquid CC2903 vs Slurry CC2903 | 0.224 | 5.54 | 0.015 |
| Between 7d Old: | Liquid Control vs Slurry Control | 0.606 | 27.7 | <0.001 |
| | Liquid Control vs Liquid CC2903 | 0.776 | 62.2 | <0.001 |
| | Liquid Control vs Slurry CC2903 | 0.721 | 46.6 | <0.001 |
| | Slurry Control vs Liquid CC2903 | 0.69 | 40.1 | <0.001 |
| | Slurry Control vs Slurry CC2903 | 0.021 | 0.386 | 0.807 |
| | Liquid CC2903 vs Slurry CC2903 | 0.78 | 71.5 | <0.001 |

Supplemental Table 3: Pairwise PERMANOVA on medfly microbiome

| Comparison | R^2 | F | p-value |
|---|---|---|---|
| Liquid Control vs Slurry Control | 0.096 | 2.23 | 0.075 |
| Liquid Control vs Liquid CC2903 | 0.457 | 20.9 | <0.001 |
| Liquid Control vs Slurry CC2903 | 0.056 | 1.26 | 0.315 |
| Slurry Control vs Liquid CC2903 | 0.389 | 11.5 | <0.001 |
| Slurry Control vs Slurry CC2903 | 0.046 | 0.673 | 0.548 |
| Liquid CC2903 vs Slurry CC2903 | 0.38 | 11.1 | <0.001 |
